# miR-1 coordinately regulates lysosomal v-ATPase and biogenesis to affect muscle contractility upon proteotoxic challenge during ageing

**DOI:** 10.1101/2021.01.21.427623

**Authors:** Isabelle Schiffer, Birgit Gerisch, Kazuto Kawamura, Raymond Laboy, Jennifer Hewitt, Martin S. Denzel, Marcelo A. Mori, Siva Vanapalli, Yidong Shen, Orsolya Symmons, Adam Antebi

## Abstract

Muscle function relies on the precise architecture of dynamic contractile elements, which must be fine-tuned to maintain motility throughout life. Muscle is also highly plastic, and remodelled in response to stress, growth, neural and metabolic inputs. The evolutionarily conserved muscle-enriched microRNA, miR-1, regulates distinct aspects of muscle biology during development, but whether it plays a role during ageing is unknown. Here we investigated the role of *C. elegans* miR-1 in muscle function in response to proteostatic stress during adulthood. *mir-1* deletion results in improved mid-life muscle motility, pharyngeal pumping, and organismal longevity under conditions of polyglutamine repeat proteotoxic challenge. We identified multiple vacuolar ATPase subunits as subject to miR-1 control, and the regulatory subunit *vha-13*/ATP6VIA as a direct target downregulated via its 3’UTR to mediate miR-1 physiology. miR-1 further regulates nuclear localization of lysosomal biogenesis factor HLH-30/TFEB and lysosomal acidification. In summary, our studies reveal that miR-1 coordinately regulates lysosomal v-ATPase and biogenesis to impact muscle function and health during ageing.

## Introduction

Ageing is the major cause of progressive decline in all organ systems throughout the body. This is particularly evident in the musculature and often manifests as sarcopenia, the loss of muscle mass and strength. In fact, muscle frailty is a hallmark of tissue ageing seen in species as diverse as worms, flies, mice and humans (Herndon et al., 2002; Miller et al., 2008) (V. G. Martinez et al., 2007) (Demontis, Piccirillo, Goldberg, & Perrimon, 2013) (Cruz-Jentoft et al., 2010) (Nair, 2005). At the molecular level, frailty is often accompanied by a decline in muscle structure and function, as well as alterations in muscle proteostasis and metabolism. Nevertheless, muscle can respond positively to exercise and stress and rejuvenate even into older age, showing remarkable plasticity (Pollock et al., 2018) (Cartee, Hepple, Bamman, & Zierath, 2016) (Distefano & Goodpaster, 2018). A molecular study of muscle ageing and plasticity in a genetically tractable model could shed light on fundamental aspects of these processes.

microRNAs are small 22-26 nucleotide RNAs that bind with complementarity through their seed sequence to target mRNAs to downregulate gene expression (Gu & Kay, 2010). They can work as molecular switches or fine tune gene regulation, and typically have multiple targets, thereby coordinating cellular programs. Many microRNAs are expressed in a tissue-specific manner and regulate programs cell autonomously, and could therefore potentially serve as tissue-specific therapeutic targets (Guo et al., 2014) (Panwar, Omenn, & Guan, 2017). In addition, some microRNAs are found circulating in serum (Weber et al., 2010) (Chen et al., 2008), raising the possibility that they can act cell non-autonomously as well.

miR-1 homologs are muscle-enriched microRNAs that are highly conserved in evolution, and important for mammalian heart and muscle development (Zhao et al., 2007). Like in other animals, *Caenorhabditis elegans mir-1* is expressed in body wall (“skeletal”) and pharyngeal (“cardiac”) muscle (Simon et al., 2008). Deletion mutants are viable and exhibit modest changes in the behavior of the neuromuscular junction (Simon et al., 2008) and autophagy (Nehammer et al., 2019), but little else is known of its normative function. In this work, we examined the potential role of miR-1 in regulating muscle function and proteostasis. Surprisingly, we found that under polyQ35 proteotoxic challenge, *mir-1* mutation prevents aggregate formation, and increases organismal motility and pharyngeal contractility during ageing. Mechanistically, miR-1 regulates expression of several lysosomal v-ATPase subunits, and targets a crucial regulatory component, *vha-13*/ATP6VIA, via its 3’ UTR to impact proteotoxicity and longevity. Further, miR-1 regulates lysosomal acidification and nuclear localization of HLH-30/TFEB, a key regulator of lysosome biogenesis. These studies reveal an unexpected role of miR-1 in coordinating lysosomal function and health during ageing.

## Results

### *mir-1* mutation improves muscle motility upon polyQ challenge

*mir-1* is a highly conserved microRNA expressed predominantly in muscle tissues including pharyngeal muscle, body wall muscle, and sex muscles (Andachi & Kohara, 2016; N. J. Martinez et al., 2008). To unravel molecular miR-1 function, we first characterized the nature of several *mir-1* mutations. The reference alleles (*n4101, n4102*) consist of large deletions of the region that could have confounding effects on nearby loci. We therefore focused on *gk276*, a smaller 192 base pair deletion that removes the *mir-1* coding region as well as the downstream non-coding region. We also generated an independent *mir-1* allele, *dh1111,* by CRISPR genome engineering (Figure 1A), which deletes a 52 base pair region within the *mir-1* locus. Both alleles failed to express the mature miRNA as measured by Taqman qPCR (Supplemental Figure 1A), and thus are *mir-1* null mutants.

**Figure 1.**
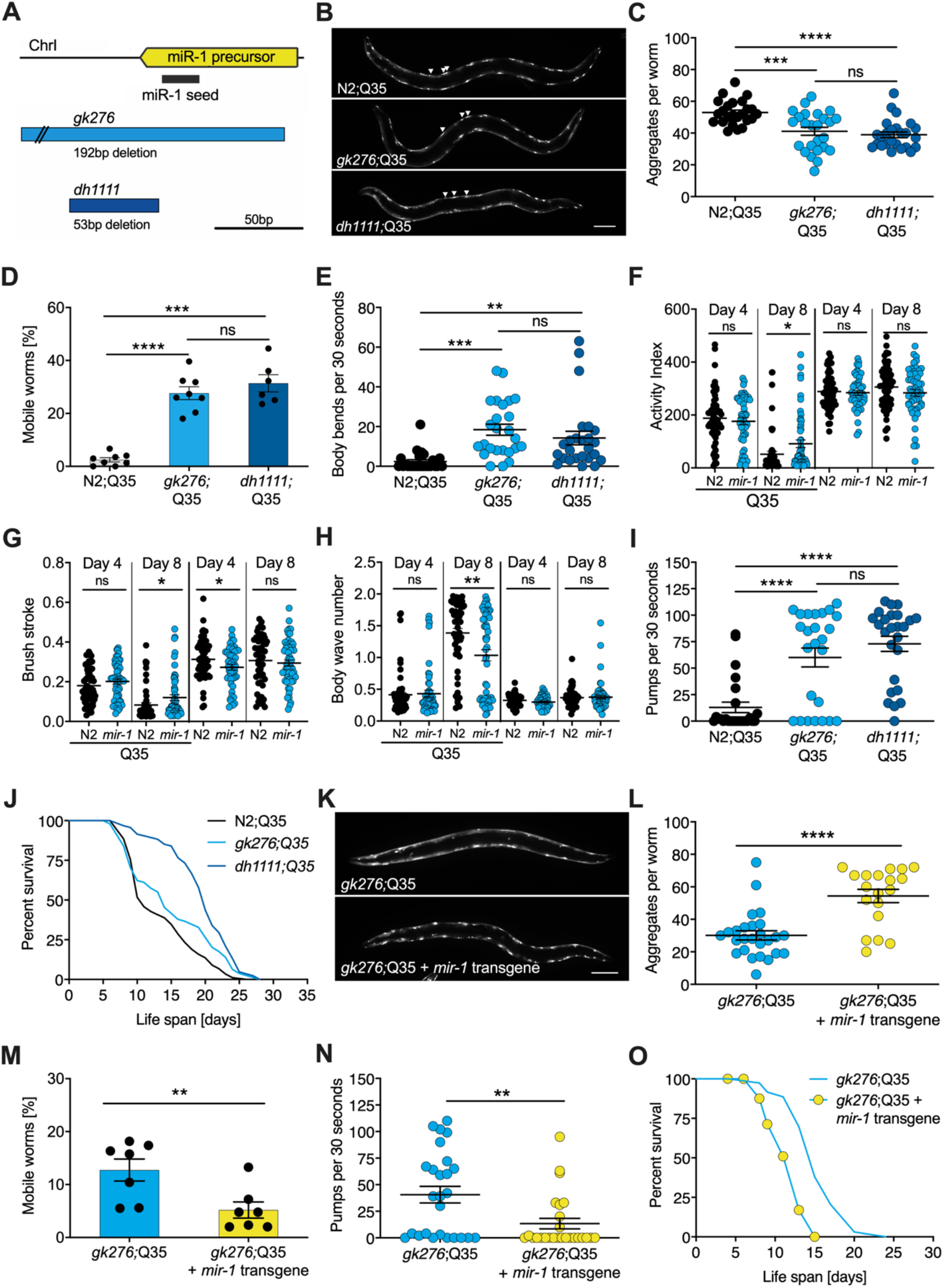
*mir-1* mutants exhibit improved motility upon polyQ challenge. **(A)** Schematic showing the *mir-1* locus, deletion alleles *mir-1(gk276)* and *mir-1(dh1111)* (see Material and Methods). **(B)** Representative images of wild-type N2 and *mir-1* mutant animals expressing *unc-54p::Q35::YFP* (Q35) showing loss of aggregates in the *mir-1* background on day 4 of adulthood. Arrowheads point to aggregates. Scale bar 100 μm. **(C)** Quantification of *Q35::YFP* aggregates in B using Zen software, each dot represents one animal, mean ± SEM from one representative experiment, N=3, one way ANOVA, Tukey′s multiple comparisons test, ***, p<0.001, ****, p<0.000, ns, not significant. **(D)** Motility of indicated *mir-1* alleles and wild-type animals expressing *unc-54p::Q35::YFP*, measured by the circle test, day 8 of adulthood. Percent worms that left the circle after 30 minutes was determined. Each dot represents one experiment. Mean ± SEM of N=6 to 8. One way ANOVA, ***, p<0.001, ****, p<0.0001, ns, not significant. **(E)** Motility of *mir-1* alleles and wild-type animals expressing *unc-54p::Q35::YFP*, measured by the thrashing assay. 25 worms per condition, each dot represents one animal, mean ± SEM from one representative experiment, N=4, one way ANOVA, **, p<0.01 ***, p<0.001, ns, not significant. CeleST analysis of activity index **(F)**, brush stroke **(G)** and body wave **(H)** comparing N2 and *mir-1(gk276)* mutant animals with or without *unc-54p::Q35::YFP* at day 4 and day 8 of adulthood. Each dot represents one animal. t-test, *, p<0.05 **, p<0.01, ns, not significant. **(I)** Pharyngeal pumping rate measured on day 8 of adulthood in *mir-1* alleles and wild-type animals expressing *unc-54p::Q35::YFP.* Each dot represents one animal, mean ± SEM from one representative experiment, N=3, one way ANOVA, ****, p<0.0001, ns, not significant. **(J)** Life span experiments of *mir-1* mutants and wild-type animals expressing *unc-54P::Q35::YFP.* One experiment of N=3. t-test: N2;Q35 *vs. gk276*;Q35: p=0.03. N2;Q35 *vs. dh1111*;Q35: p<0.0001. **(K)** Presence of an extrachromosomal *mir-1* transgene brings back aggregates in *mir-1(gk276)* mutants at day 4 of adulthood, showing rescue by the transgene. Transgenic worms were compared to non-transgenic segregants of the same strain. Scale bar 100 μm. **(L)** Quantification of *Q35::YFP* aggregates (from **K**), each dot represents total aggregate number of one animal, mean ± SEM from one representative experiment, N=3, t-test, ****, p<0.0001. **(M)** Motility of *mir-1(gk276) unc-54p::Q35::YFP* animals in the presence or absence of the *mir-1* transgene at day 8 of adulthood, measured by circle test. Each dot represents one experiment, mean ± SEM of N=7. Total number of worms tested 646 (transgenic worms) and 469 (non-transgenic segregants of the same strain), t-test, **, p<0.01. **(N)** Pharyngeal pumping rate of *mir-1(gk276) unc-54p::Q35::YFP* animals in the presence or absence of the *mir-1* transgene. Transgenic worms were compared to non-transgenic segregants of the same strain. Each dot represents one animal, mean ± SEM from one representative experiment, N=7, t-test,**, p<0.01. **(O)** Life span experiments of *mir-1(gk276) unc-54p::Q35::YFP* animals in the presence or absence of the *mir-1* transgene. Transgenic worms were compared to non-transgenic segregants of the same strain. One representative experiment of N=5, t-test, p<0.0001.

To investigate *mir-1* function, we measured several physiological parameters. We found that *mir-1* mutants *gk276* and *dh1111* developed normally, had normal brood size, and near normal life span (Supplemental Figure 1B-D, Supplemental Table 1). Additionally, motility on solid media, thrashing in liquid culture, and pharyngeal pumping ability determined at day 8 and 14 of adulthood were similar to wild-type N2 controls (Supplemental Figure 1E-G, Supplemental Table 2). Since *mir-1* deletion showed little obvious phenotype on its own, we reasoned that some physiological differences might emerge under stress. However, we saw little distinction from wild-type (WT) with moderate heat stress by growth at 25°C nor heat shock resistance at 35°C (Supplemental Figure 1D and H, Supplemental Table 1).

We next tested the idea that *mir-1* mutants might differ in their response to proteotoxic stress. To do so, we used a strain expressing polyglutamine-35 tracts fused to yellow fluorescent protein (YFP) under the control of the muscle myosin specific *unc-54* promoter (*unc-54p::Q35::YFP*), which has been used previously in models of proteotoxicity and Huntington’s disease (Morley, Brignull, Weyers, & Morimoto, 2002) (Brignull, Morley, Garcia, & Morimoto, 2006). In these strains, initially soluble proteins become sequestered into insoluble aggregates, visible as foci in the muscle of the worm. Worms expressing the polyQ35 stretches showed progressive age-dependent protein aggregation, which became clearly visible by day 4 of adulthood. Interestingly, we observed that *mir-1* mutants displayed significantly less aggregates (avg. 38 and 36) compared to age-matched wild-type controls (avg. 53) at this time (Figure 1B and C, Supplemental Table 2).

We further investigated the possible effect of proteotoxicity by measuring organismal motility of Q35 strains using the circle test. This test measures how many worms crawl out of a 1 cm circle within 30 minutes on day 8 of adulthood on agar plates. Day 8 was chosen as a timepoint where paralysis becomes visible but animals are still viable. We observed that about 30% of the *mir-1;*Q35 mutants left the circle while only 2.5% of the control Q35 worms did so (Figure 1D, Supplemental Table 2). We also measured motility by counting the thrashing rate in liquid culture on day 8 of adulthood. Though the degree of paralysis was more variable, both *mir-1* mutants *gk276* and *dh1111* were significantly more mobile (average 16, 15 body bends) compared to age-matched wild-type controls (average 2 body bends, Figure 1E, Supplemental Table 2).

To further characterize miR-1’s impact on muscle function, we performed CeleST analysis (Restif et al., 2014), a MATLAB based algorithm that extracts various parameters of movement (wave initiation, travel speed, brush stroke, activity index) and gait (wave number, asymmetry, curl, stretch) from video footage. Typically activity measures decline with age, whereas gait abnormalities increase with age (Restif et al., 2014). We scored animals focusing on day 4 and 8 of adulthood comparing *mir-1* and WT, in the presence and absence of the Q35 array. In the absence of Q35, *mir-1* mutants performed similar to WT (*e.g.*, body wave) or were slightly different (wave initiation, travel speed, stretch) for some parameters (Supplemental Table 3). In the presence of Q35, however, *mir-1* mutants did significantly better than WT for several measures of activity (activity index, brush stroke, travel speed, wave initiation) and gait (body wave) on day 8 (Figure 1F-H, Supplemental Figure 1I-M, Supplemental Table 3), consistent with the idea that *mir-1* mutation protects against proteotoxic stress during mid-life.

*mir-1* is also highly expressed in the pharynx, a contractile tissue similar to cardiac muscle. We observed that in the *mir-1*;Q35 background, pumping rate was significantly increased (*gk276*, avg. 60; *dh1111*, avg. 78) compared to Q35 controls (avg. 16) on day 8 of adulthood (Figure 1l, Supplemental Table 2). Furthermore, we found that the life span of *mir-1(dh1111)*;Q35 strains was significantly extended (N=3, mean 13 (WT) vs.19 (*dh1111*) days) compared to Q35 alone (Figure 1J, Supplemental Table 1). A similar trend was seen with *gk276* in 2 of 3 experiments (Figure 1J, Supplemental Table 1). These findings suggest that *mir-1* reduction can counter the intrinsic and systemic detrimental effects of Q35 proteotoxic challenge.

Introducing a wild-type *mir-1* transgene into the *mir-1(gk276)* background restored normal levels of *mir-1* expression (Supplemental Figure 1N) and largely reversed *mir-1* phenotypes, showing increased Q35 aggregates, decreased motility, pumping rate, and life span compared to non-transgenic controls (Figure 1K-O, Supplemental Table 1 and 2). The similarity of behaviour among the different *mir-1* alleles and the rescue of these phenotypes by the wild-type transgene demonstrate that the *mir-1* mutation is causal for removing muscle aggregates and improving mid-life motility.

### v-ATPase subunits are downstream mediators of miR-1 induced motility improvement

To identify downstream mediators of protein quality control improvement in *mir-1* mutants, we used computational and proteomic approaches to find potential miR-1 targets. For the computational approach, we used publicly available prediction tools to identify genes harbouring miR-1 seed binding sites in their 3’UTR, namely microRNA.org, TargetScanWorm, and PicTar (Betel, Wilson, Gabow, Marks, & Sander, 2008) (Jan, Friedman, Ruby, & Bartel, 2011) (Lall et al., 2006). These prediction tools use distinct algorithms that often yield different and extensive sets of candidates (Figure 2A). Therefore, we limited ourselves mostly to candidates that were predicted as targets by all three algorithms. This yielded 68 overlapping candidates (Figure 2A), 50 of which had available RNAi clones (Supplementary Table 4). Interestingly, network analysis of these 68 genes using STRING (string-db.org) (Snel, Lehmann, Bork, & Huynen, 2000) revealed a tight cluster of predicted targets encoding 11/21 subunits of the vacuolar ATPase (Figure 2B, Supplemental Figure S2A). 5 additional vacuolar ATPase genes were identified by 2 of the 3 prediction algorithms (Supplemental Figure 2A). In addition, two genes that code transcription factors implicated in lysosome biogenesis, *daf-16*/FOXO and *hlh-30*/TFEB (Sardiello et al., 2009; Settembre et al., 2011), also contained predicted miR-1 binding sites in their 3’UTRs.

**Figure 2.**
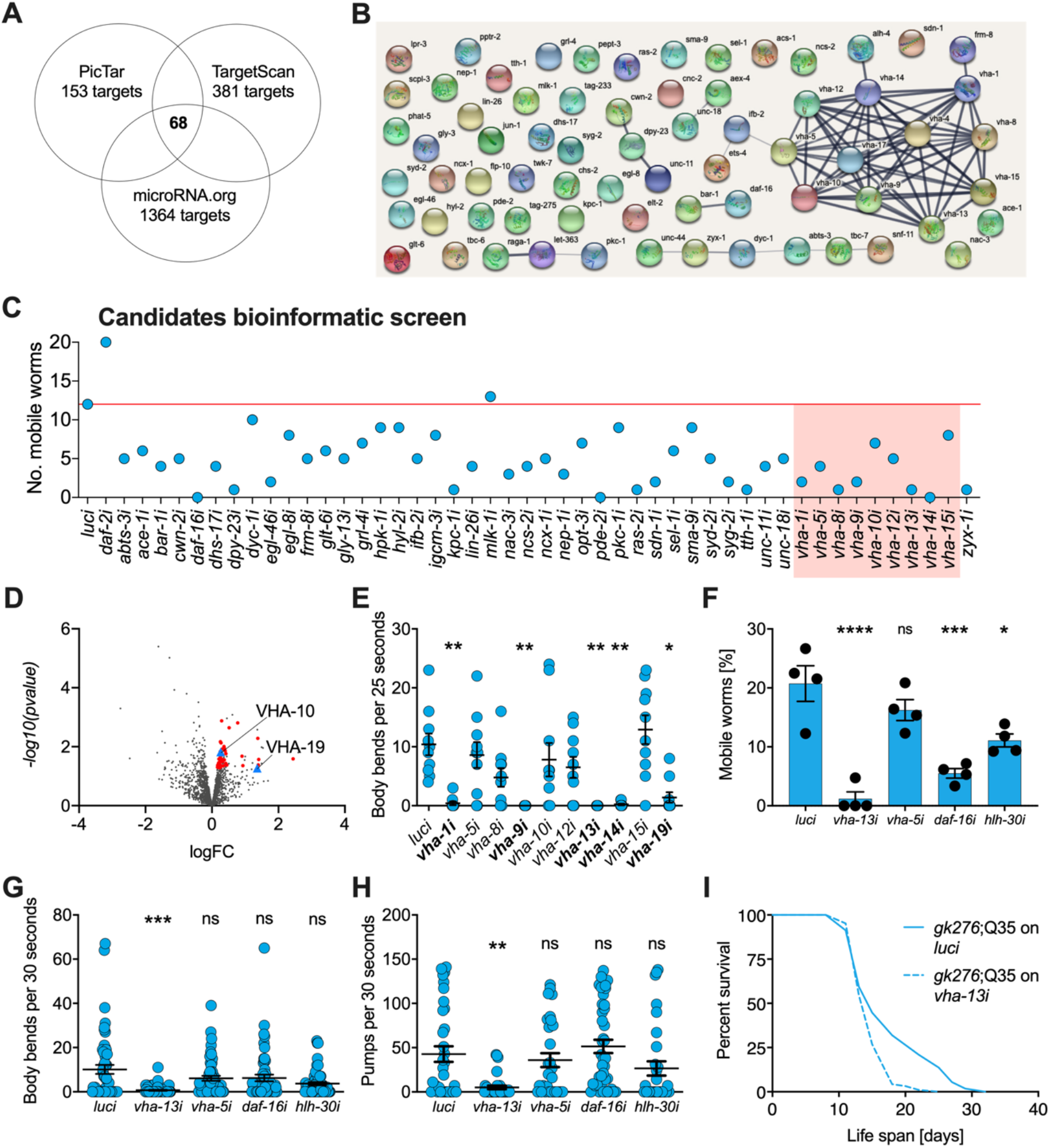
v-ATPase subunits are downstream mediators of *mir-1* induced motility improvement. **(A)** Computational screen using *in silico* prediction of miR-1 binding sites in target mRNAs using microRNA.org, TargetScan and PicTar identifies 68 shared candidates. **(B)** STRING network analysis of predicted miR-1 targets reveals a cluster of vATPase subunits. **(C)** Initial RNAi screen of computationally predicted candidates using the circle test on day 8 of adulthood reveals a number of candidates that reduced motility of *mir-1(gk276)*;Q35, 20 worms per RNAi (N=1). Red line, number of luciferase controls that left the circle. v-ATPases are highlighted in red. **(D)** Volcano plot of proteomic analysis showing differentially regulated proteins in *mir-1* vs. N2 animals. Red dots show upregulated proteins tested in circle assay, VHA-10 and VHA-19 indicated by blue triangles. N=6 (*mir-1*) and 4 (N2) biological replicates. **(E)** Effect of v-ATPase subunit RNAi knockdown on *mir-1(gk276)*;Q35 motility as measured in the thrashing assay. Animals are grown on the corresponding RNAi from L4 onwards. N=1. Mean ± SEM, one way ANOVA, only significant values are labelled: *, p<0.05, **, p<0.01. **(F)** and **(G)** Motility assay (circle test) and trashing assay on day 8 of *mir-1(gk276)*;Q35 worms upon *vha-13, vha-5, daf-16 and hlh-30* RNAi knockdown. Control, luciferase RNAi (*luci*). Mean ± SEM of N=4, one way ANOVA, *, p<0.05, ***, p<0.001, ****, p<0.0001, ns, not significant. **(H)** Pumping assay of *mir-1(gk276)*;Q35 worms upon *vha-13, vha-5, daf-16 and hlh-30* RNAi knockdown on day 8 of adulthood. Control, luciferase RNAi (*luci*). N=3, mean ± SEM, one way ANOVA, **, p<0.01, ns, not significant. **(I)** Life span of *mir-1*;Q35 and Q35 worms upon *vha-13* RNAi knockdown. One experiment of N=3, t-test: p<0.01.

Next, we carried out a functional screen of the selected candidates. We reasoned that *mir-1* deletion likely results in an upregulation of these proteins, and that RNAi knockdown of the relevant genes should therefore abrogate the improved motility of *mir-1*;Q35 worms. Specifically, we scored motility using a rapid version of the circle test, measuring the ability of *mir-1*;Q35 worms grown on candidate RNAis to migrate out of a 1 cm diameter circle within 1 minute (Figure 2C). 12/20 *mir-1*;Q35 worms left the circle when grown on luciferase control RNAi (*luci)*. We considered genes as potential *mir-1*;Q35 suppressors if less then 4 worms left the circle upon RNAi knock down of the genes, and prioritized candidates when only 0 or 1 worms left. Among the selected candidates vacuolar ATPase subunits *vha-1, vha-8, vha-9, vha-13,* and *vha-14*, as well as muscle gene *zyx-1*, stood out as molecules suppressing *mir-1* motility (Figure 2C and Supplemental Figure 2D).

As a second parallel approach to identify miR-1 regulatory targets in an unbiased manner, we performed TMT shotgun proteomic analysis, comparing *mir-1* mutants to WT worms on day 1 of adulthood. We identified approximately 1,500-2,000 proteins in the replicates (Figure 2D). Samples showed a high degree of correlation (Supplemental Figure 2B) with minimal separation of WT and *mir-1* genotypes, reflecting the subtle changes induced by *mir-1* mutation at the organismal level. For further analysis we therefore selected the 50 top upregulated proteins with available RNAi clones and tested them for their ability to suppress the motility phenotype using the circle test (Supplemental Figure 2C and 2E, Supplemental Table 5), among them *vha-10*, *vha-19* and *zyx-1*, which had previously been predicted to harbour *mir-1* binding sites.

Based on the prominent presence of vATPase genes among both the informatic and the proteomic candidates, we systematically tested 10 of them in the thrashing assay, confirming v-ATPase subunits *vha-1, vha-9, vha-13, vha-14* and *vha-19* as likely mediators of *mir-1*;Q35 motility (Figure 2E). We further tested the impact of *vha-5, vha-13* and transcription factors *daf-16/FOXO* and *hlh-30/TFEB* RNAi *on mir-1*;Q35 phenotypes in more detail, using the circle test, thrashing assay and pharyngeal pumping as readout (Figure 2F-H), which again confirmed in particular the role of *vha-13* in reducing *mir-1* motility. We also confirmed that the effect on motility was specific to the *mir-1* background for *vha-5*, *vha-10*, *vha-13* and *hlh-30*, since RNAi of these genes had no significant effect on motility in N2;Q35 worms (Supplemental Figure 2F).

In summary, knockdown of 5 of 10 tested v-ATPase subunits from both the computational and proteomics screen, as well as 2 transcriptional mediators of lysosome biogenesis, reduced or abolished *mir-1* motility improvement either by circle test or thrashing, revealing these molecules to have congruent physiological effects. Therefore, we focused most of our further efforts on examining v-ATPase regulation by miR-1.

### miR-1 directly regulates VHA-13 protein levels in muscle tissue

MicroRNAs regulate their targets by binding to the 3’ UTR of client mRNAs, thereby decreasing RNA stability and translation. If miR-1 regulates v-ATPase subunits, then miR-1 loss would be predicted to de-repress v-ATPase mRNA and protein levels. Because the onset of Q35 aggregation becomes visible by day 4 of adulthood, we examined the mRNA expression of representative v-ATPase subunits at this time. We found that the mRNA levels of *vha-5, vha-10, vha-13*, and *vha-19,* as well as transcription factors *hlh-30*/TFEB and *daf-16*/FOXO were increased approximately 2-fold in *mir-1* mutants compared to age-matched wild-type worms (Figure 3A), consistent with the idea that these genes are miR-1 targets.

**Figure 3.**
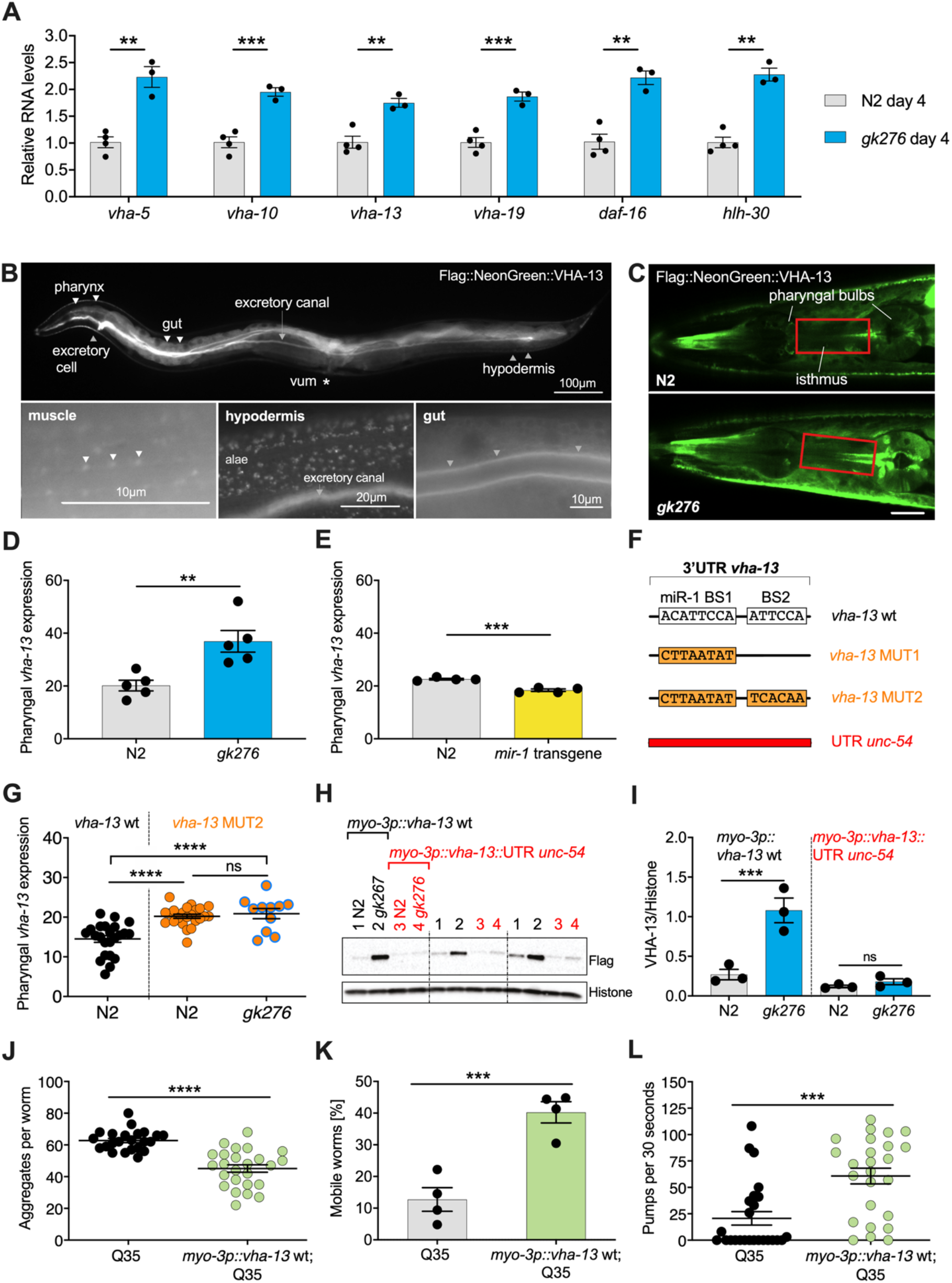
miR-1 directly regulates *vha-13* via its 3’UTR in muscle tissue. **(A)** RT-qPCR of *vha-5, vha-10, vha-13* and *vha-19*, as well as *daf-16* and *hlh-30* mRNA levels in WT (N2) and *mir-1(gk276)* mutants on day 4 of adulthood. Mean ± SEM, N=3, one way ANOVA, **, p<0.01, ***, p<0.001. **(B)** Expression pattern of endogenously tagged *3xFlag::mNeonGreen::vha-13* in pharynx, excretory cell and canal, gut, vulva muscle (vum), hypodermis, muscle (arrowhead indicates dense body). **(C)** Confocal images of the head region in worms carrying endogenously tagged *3xFlag::mNeonGreen::vha-13* in late L4 *mir-1* and WT animals at 25°C. Red rectangle highlights the area of *vha-13* expression in the isthmus used for determination of mNeonGreen intensity. **(D)** Quantification of fluorescent intensity of *3xFlag::mNeonGreen::vha-13* expression in the isthmus of indicated genotypes (as shown in **(C)**) in late L4 larvae at 25°C. Mean ± SEM of N=5, t-test, **, p<0.01. **(E)** Quantification of fluorescent intensity of *3xFlag::mNeonGreen::vha-13* expression in the isthmus of N2; *mir-1* transgenic overexpression strain in late L4 larvae at 25°C. Transgenic worms (*mir-1* transgene) were compared to non-transgenic segregants (N2) of the same strain. Mean ± SEM of N=4, t-test, ***, p<0.001. **(F)** Graphic showing the 3’UTR of endogenously tagged *3xFlag::mNeonGreen::vha-13. vha-13 wt: vha-13* wild-type 3′UTR. *vha-13* MUT1: one miR-1 binding site (BS) is mutated. *vha-13* MUT2: both miR-1 BSs are mutated. *unc-54*: the *vha-13* 3’UTR is substituted by *unc-54* 3’UTR. Nucleotide sequence of mutated miR-1 binding sites are shown below WT. **(G)** Quantification of fluorescence intensity in the isthmus of L4 larvae with endogenously tagged *3xFlag::mNeonGreen::vha-13 vha-13 MUT2* 3’UTR in relation to *vha-13 wt* 3’UTR, in N2 and *mir-1(gk276)* mutant backgrounds at 25°C, using confocal imaging. Mean ± SEM of N=3, one representative experiment, one-way ANOVA, ****, p<0.0001, ns not significant. **(H)** Western blot image of late L4 WT and *mir-1* mutants expressing transgenic flag-tagged *vha-13* in body wall muscle (*myo-3p*::*vha-13*) with either the *unc-54* 3′UTR (labelled in red), which lacks miR-1 BSs, or the wt *vha-13* 3′UTR which contains the two miR-1 BSs (labelled in black), immunoblotted with anti-Flag and anti-Histone H3 antibodies. Histone H3 loading control is shown below. Biological replicates (N=3) separated by dashed lines. **(I)** Quantification of Western blot shown in **(H)**, with flag-tag intensity normalized to histone H3 loading control, mean ± SEM of N=3, one-way ANOVA, ***, p<0.001, ns, not significant. **(J)** Quantification of aggregates in Q35 worms expressing transgenic *vha-13* in the body wall muscle (*myo-3p*::*vha-13*;Q35) or non-transgenic segregants (Q35) of the same strain. 25 worms per genotype, mean ± SEM of one representative experiment, N=4, t-test, ***, p<0.001. **(K)** Motility of Q35 worms expressing transgenic *vha-13* in the body wall muscle (*myo-3p*::*vha-13*;Q35) or non-transgenic segregants (Q35) of the same strain in circle test. Mean ± SEM. One experiment of N=2, t-test, ***, p<0.001. **(L)** Pharyngeal pumping rate of *myo-3p*::*vha-13*;Q35 and Q35 non-transgenic segregants of the same strain. One representative experiment of N=4, mean ± SEM, t-test, ***, p<0.001.

To investigate whether miR-1 also regulates candidates at protein level, we endogenously tagged the v-ATPase subunits *vha-5, vha-10, vha-13* and *vha-19* by CRISPR/Cas9 using an N-terminal 3xFlag-mNeonGreen-tag. Tagging *vha-19* and *vha-10* caused lethality and were not pursued further. Tagged *vha-5* showed significant upregulation in the pharynx in *mir-1* mutants compared to wild-type, but not in vulva muscle (Supplementary Figure 3A and B, Supplemental Table 7). Furthermore, RNAi knockdown only weakly suppressed *mir-1* motility (Figure 2E-G, Supplemental Table S2). N-terminally tagged *vha-13(syb586),* however, was viable, showed expression that was regulated, and harboured two potential *mir-1* binding sites in the 3’ UTR (Figure 3F, Supplemental Figure 3C). Moreover, *vha-13* RNAi (*vha-13i)* significantly compromised the motility of *mir-1*;Q35 strains in the RNAi screens (Figure 2C), which we further confirmed through thrashing assays and the circle test (Figure 2E-G). *vha-13i* also significantly reduced pharyngeal pumping and life span in the *mir-1*;Q35 background compared to *luci* RNAi controls (Figures 2H and I, Supplemental Tables 1 and 2), yet had relatively minor effects on the motility of N2;Q35 itself (Supplemental Figure 2F). We therefore focused on *vha-13* as a promising candidate to pursue in depth for regulatory interactions with miR-1.

We first characterized the expression pattern of *3xFlag::mNeonGreen::vha-13(syb586)* in more detail. The fusion protein was strongly expressed in the excretory cell, canal, hypodermis, as well as gut (Figure 3B). In the hypodermis *3xFlag::mNeonGreen::VHA-13* was localized in discrete foci (Figure 3B, Supplemental Figure 3D). Notably, *3xFlag::mNeonGreen::VHA-13* also resided in miR-1 expressing tissues of body wall muscle, sex muscles and pharynx, where it was more weakly expressed. Within muscle cells, VHA-13 localized to dense bodies (analogous to vertebrate Z disks) and intervening cytosol (Figure 3B, Supplemental Figure 3D).

Because *mir-1* is expressed in muscle tissues, we focused on quantifying the expression of *vha-13* in these tissues. Strikingly, we observed that *VHA-13* levels were upregulated in the pharyngeal isthmus of *mir-1* mutants (Figure 3C and D, Supplemental Table 7), while conversely, overexpression of the *mir-1* transgene decreased levels in this tissue relative to wild-type (Figure 3E, Supplemental Table 7). To further investigate whether miR-1 directly regulates *vha-13*, we altered two predicted miR-1 binding sites (BS) in the 3’UTR (Figure 3F). We first mutated miR-1 binding site 1 (*vha-13* BS1 *(syb504)*, at position 188-195 of *vha-13* 3’UTR), and then additionally miR-1 binding site 2 (*vha-13* BS2 *(syb2180)*, at position 253-58 of *vha-13* 3’UTR) in the endogenously-tagged *vha-13(syb586)* strain. When only BS1 was mutated, we still saw residual regulation of VHA-13 by miR-1 (Supplemental Figure 3E, Supplemental Table 7). Mutating both miR-1 binding sites (BS1,2: MUT2), however, de-repressed VHA-13 levels in the isthmus in the WT background (Figure 3G). In double mutants containing both *mir-1* and *vha-13* BS1,2 mutations, no clear regulation of *vha-13* expression was observed compared to the BS1,2 mutations alone (Figure 3G), suggesting that miR-1 modulates pharyngeal *vha-13* expression through both potential miR-1 BS sites.

We next sought to characterize *vha-13* regulation in the body wall muscle. However, it was not possible to accurately measure expression in this tissue using the *3xFlag::mNeonGreen::vha-13* strain due to the high expression levels in the adjacent hypodermis and gut. We therefore generated transgenic worms expressing flag-tagged *vha-13* containing its endogenous 3′UTR *(3xFlag::vha-13::vha-13* 3′UTR) under control of the muscle-specific *myo-3* promoter. Consistent with results seen in pharyngeal muscle, *vha-13* protein levels in the body wall muscle were also upregulated in *mir-1* mutants compared to WT, as measured by Western blots (Figure 3H and I). Exchanging the *vha-13* 3′UTR with the *unc-54* 3’UTR lacking miR-1 binding sites (*vha-13::unc-54 3′UTR*) blunted regulation by miR-1 (Figure 3H and I), indicating that *vha-13* is repressed by miR-1 via its 3’UTR in both body wall and pharyngeal muscle.

Since *mir-1* mutation resulted in *vha-13* upregulation, we asked whether *vha-13* overexpression was sufficient to yield phenotypes similar to *mir-1* mutants. In accord with this idea, overexpressing *vha-13* in the muscle of Q35 worms significantly reduced aggregate number, improved motility, and enhanced pharyngeal pumping ability (Figure 3J-L, Supplemental Figure 3F, Supplemental Table 2). It also significantly increased the life span of Q35 animals compared to non-transgenic controls in 2/4 experiments (Supplemental Figure G, Supplemental Table 1). Altogether these data suggest that miR-1 and VHA-13 work in the same regulatory pathway to influence muscle physiology.

### miR-1 regulates *hlh-30*/TFEB transcription factor

Our RT-PCR experiments indicated that miR-1 also regulates mRNA levels *hlh-30/TFEB* and *daf-16*/FOXO transcription factors (Figure 3A). These factors modestly affected motility in the circle test (Figure 2F), and we wondered if regulation extended to the protein level. When we examined endogenously tagged *hlh-30*, we saw no obvious regulation of protein expression levels when measured by fluorescence intensity in the pharynx (Supplemental Figure 4A and B, Supplemental Table 7). Surprisingly, however, we observed that nuclear localization of HLH-30::mNeonGreen in the hypodermis of day 4 adults grown at 25°C was significantly increased in the *mir-1* background (Figure 4A and 4B). This finding suggests that miR-1, either directly or indirectly, regulates HLH-30 in distal tissues.

**Figure 4.**
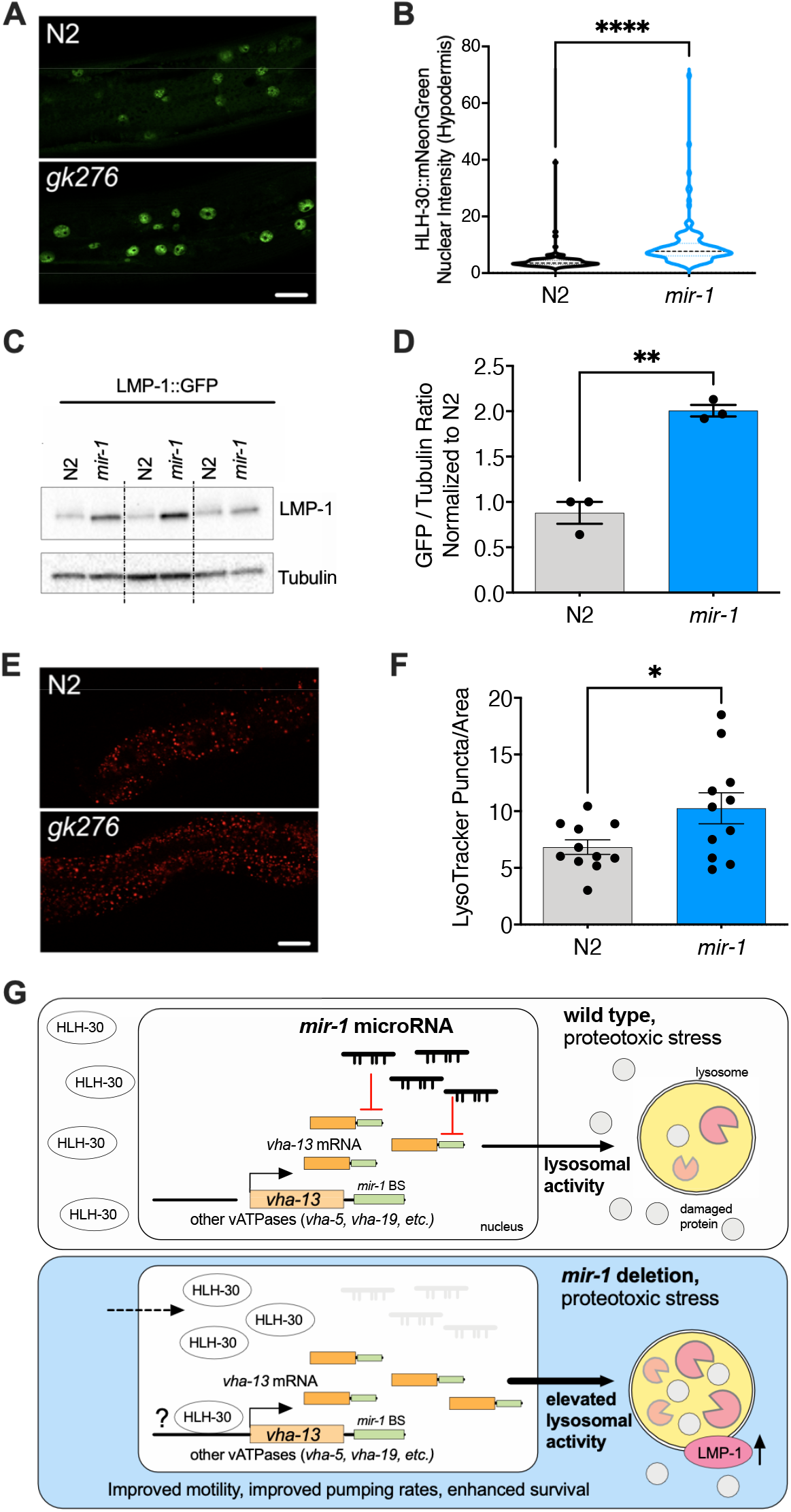
*mir-1* mutation enhances lysosomal biogenesis and acidification. **(A)** Fluorescent image comparing HLH-30::mNeonGreen nuclear localization in the hypodermis of the *mir-1(gk276)* and WT (N2) backgrounds. **(B)** Quantitation of nuclear localization in (A). Violin plot, mean ± SEM of one representative experiment of N=3, t-test, ****, p<0.0001. **(C)** Western blot of LMP-1::GFP in WT and *mir-1* mutants at the L4 stage, immunoblotted with anti-GFP or anti-α-tubulin antibodies. Biological replicates (N=3) separated by dashed lines. **(D)** Quantification of the Western blot in **(C),** normalized to α-tubulin loading control. N=3 BR, line and error bars indicate mean ± SEM, t-test, **, p<0.01. **(E)** Representative images of lysotracker staining in WT and *mir-1* mutants at day 4 of adulthood at 25°C. **(F)** Quantification of lysotracker images using a predefined squared area approximately spanning the second to fourth gut cell. Quantification was performed using Image J. N=3 BR, line and error bars indicate mean ± SEM of combined experiments, t-test, *, p<0.05. **(G)** Working model. miR-1 normally limits proteoprotective pathways, downregulating the expression of *vha-13,* other v-ATPases and factors by binding miR-1 binding site(s) in the 3’UTR (miR-1 BS) of the corresponding mRNA (WT, proteotoxic stress). Loss of miR-1 under proteotoxic conditions results in free *vha-13* mRNA, higher VAH-13 protein levels, and elevated lysosomal activity (LMP-1). Nuclear localization of HLH-30/TFEB, a master regulator of lysosome biogenesis is also enhanced, collectively resulting in improved mid-life muscle motility, pharyngeal pumping, and organismal longevity under proteotoxic stress conditions.

### miR-1 affects lysosomal biogenesis

As the v-ATPase is an integral component of the lysosome, we asked wether miR-1 generally affects lysosomal structure and function. To test this idea, we first used a *lmp-1p::lmp-1::gfp* reporter strain. LMP-1 protein localizes to the membrane of lysosomes and is a marker for lysosomal biogenesis (Hermann et al., 2005). LMP-1 protein levels were significantly increased in *mir-1* mutants compared to WT control, as measured by Western blot analysis (Figure 4C and D). The v-ATPase hydrolyses ATP to pump protons across the membrane, resulting in acidification of the lysosome lumen (Beyenbach & Wieczorek, 2006). We therefore asked whether *mir-1* mutants affect the number of acidified lysosomes. Using Lysotracker Red, a dye that targets acidic membranous structures such as lysosomes (Chazotte, 2011), we observed an increase in the number of acidified puncta in *mir-1* mutants at day 4 of adulthood (Figure 4E and F). Due to technical limitations, lysotracker staining could only be observed in the worm intestine. This result leaves open the question as to whether lysosomes are also regulated in the muscle and whether *mir-1*, being expressed in muscle tissue, might have a cell non-autonomous effect. Nonetheless, the overall increase in lysosome biogenesis is consistent with the upregulation of lysosomal components, such as the v-ATPase subunits, and a possible cause of miR-1-dependent regulation of proteostasis.

## Discussion

Striated muscle is one of the most highly ordered tissues in the body, with molecular components organized in a lattice of contractile elements and attachments. This molecular apparatus is exposed to high energy and force during contraction, inflicting molecular damage requiring constant repair. Further, muscle is subject to growth, metabolic, and stress signalling pathways as well as neural inputs that also promote remodelling. Underlying muscle plasticity is the fine-tuned control of proteostasis, including protein synthesis, folding, trafficking and turnover, which must be precisely orchestrated to maintain muscle structure and function (Demontis et al., 2013). A decline in these processes during aging leads to diminished muscle performance and frailty, yet it remains elusive how various aspects of muscle proteostasis are coordinated.

In this work, we discovered that the muscle enriched microRNA miR-1 plays an important role in regulating muscle homeostasis via vacuolar ATPase function. Loss of *mir-1* ameliorates age-related decline in motility induced by models of aggregate-prone polyQ35. By inference, *mir-1* normally limits proteoprotective pathways. Computational and proteomic screens identified v-ATPase subunits as highly enriched targets of miR-1 regulation, suggestive of coordinate regulation, and whose downregulation reduced proteoprotection. Expression studies confirmed that several subunits (*e.g*., *vha-5, vha-10, vha-13, vha-19*) showed miR-1 dependent regulation. In particular, we demonstrated that *vha-13* expression is downregulated by miR-1 in muscle tissues via two distinct binding sites in its 3’UTR. Moreover, VHA-13 links miR-1 with muscle homeostasis, since *vha-13* downregulation abolished the improved mid-life motility, pharyngeal pumping and life span of *mir-1*;Q35 strains, while *vha-13* overexpression was sufficient to enhance these properties, similar to *mir-1* mutation. In accord with our findings, immunoprecipitation and sequencing of microRNA complexes revealed a number of v-ATPase subunit mRNAs, including *vha-13*, *vha-4, vha-10,* and *vha-17* as physically associated with *C. elegans* miR-1 (Grosswendt et al., 2014). Interestingly, mammalian homolog of VHA-13, ATP6V1A, as well as several other v-ATPase subunits (Stark, Brennecke, Bushati, Russell, & Cohen, 2005), harbour predicted miR-1 binding sites, suggesting that this regulatory module could be conserved in evolution.

The v-ATPase is a multisubunit enzyme which acidifies the endolysosomal lumen to control a plethora of cellular activities. Acidification regulates protein trafficking, endocytic recycling, synaptic vesicle loading, and autophagy, as well as the activity of multiple acid hydrolases and nutrient and ion transporters. Moreover, the v-ATPase itself serves as a docking site for regulating mTOR and AMPK complexes and affects metabolism (Settembre, Fraldi, Medina, & Ballabio, 2013; Zhang et al., 2014). Our studies highlight the importance of v-ATPase activity to muscle performance. A handful of studies ascribe a role for the v-ATPase in muscle. In mammalian cardiomyocytes, lipid loading inhibits the v-ATPase, leading to a decline in contractile function that could contribute to muscle deficits in diabetes (Wang et al., 2020). Lesions in VMA21 that disrupt v-ATPase assembly have also been shown to cause myopathies (Dowling, Moore, Kalimo, & Minassian, 2015). Conceivably the v-ATPase could play an important role in protein turnover and remodeling of muscle structure but could also impact muscle homeostasis through metabolic wiring or protein trafficking.

Surprisingly we also found that some *mir-1* phenotypes were not strictly limited to muscle, since we observed a global increase in levels of lysosomal LMP-1::GFP in Western blots, increased lysosomal acidification in the intestine, as well as enhanced nuclear localization of HLH-30/TFEB in the hypodermis. Given that *mir-1* is muscle-expressed, this observation could suggest cell non-autonomous regulation of lysosomal biogenesis and associated activities by *mir-1*, acting either directly or indirectly. Conceivably, miR-1 is secreted from muscle to affect physiology in other tissues. In mammals miR-1 has been identified as a circulating microRNA found in serum exosomes upon exercise stress or cardiac infarction, suggesting it could act systemically (Cheng et al., 2019) (D’Souza et al., 2018). In this light, it is intriguing that the v-ATPase itself is implicated in exosomal activity activity (Liegeois, Benedetto, Garnier, Schwab, & Labouesse, 2006). Alternately, *mir-1* could act indirectly through production of a myokine that affects distal tissues.

miR-1 has been implicated in regulating a number of targets and physiologic processes. In mammals, miR-1 and its homolog miR-133 are essential to cardiac development and function, where they have been shown to regulate Notch ligand Dll-1, the GTPase dynamin 2 (DNM2), FZD7 (Frizzled-7), and FRS2 (fibroblast growth factor receptor substrate 2) as targets (Ivey et al., 2008; Liu et al., 2011; Mitchelson & Qin, 2015). Surprisingly knockout of mammalian *mir1/mir133* specifically in muscle has little overt effect on muscle structure, but rather regulates the developmental transition from glycolytic to oxidative metabolism via the MEF-2/Dlk1-Dio3 axis, affecting running endurance (Wust et al., 2018). Similarly, *C. elegans* miR-1 regulates *mef-2*, in this case affecting retrograde signalling at the neuromuscular junction (Simon et al., 2008). We also observed that animals lacking *mir-1* have little overt change in muscle structure or function alone, though we saw upregulation of a number of muscle proteins and downregulation of mitochondrial proteins in our proteomic analysis. Indeed, most microRNA knockouts do not lead to observable phenotypes in *C. elegans* (Miska et al., 2007), suggesting that microRNAs often work redundantly or in response to stress to fine tune gene expression.

Recently Pocock and colleagues reported that miR-1 downregulates the rab GTPAse TBC-7/TBC1D15, thereby promoting autophagic flux in worms and human cells (Nehammer et al., 2019). In *C. elegans* they observed that *mir-1(+)* ameliorates polyQ40 proteotoxicity, seemingly contradicting our results. Other studies suggest that mammalian *mir-1* can either promote or inhibit autophagy, dependent on context (Ejlerskov, Rubinsztein, & Pocock, 2020; Hua, Zhu, & Wei, 2018; Xu, Cao, Zhang, Zhang, & Yue, 2020). Whether miR-1 is proteoprotective or limits proteoprotection in *C. elegans* could hinge on many factors. We used polyQ35, Pocock et al used polyQ40; these species differ in the effect on aggregation and motility (Morley et al., 2002). For motility assays, we assessed all animals that were alive, they excluded paralyzed animals. We performed motility assays on day 8, they performed them earlier in adulthood. Because microRNAs generally fine tune regulation, and often work in feedback circuits, their impact on physiology can be subtle, and small differences in culture conditions, genetic background, assay method, or metabolic state could give rise to divergent outcomes.

Among the putative miR-1 targets that we identified in *C. elegans* are the pro-longevity transcription factors DAF-16/FOXO and HLH-30/TFEB, which regulate lysosome biogenesis, proteostasis, and metabolism, and act in insulin/IGF and mTOR signalling pathways (Lin et al., 2018; O’Rourke & Ruvkun, 2013; Settembre et al., 2011). *mir-1* mutation led to an upregulation of their mRNA during adulthood and enhanced nuclear localization of HLH-30. Further RNAi knockdown of these factors modestly diminished the motility improvement of *mir-1* mutants in the circle test, consistent with roles in a miR-1 signalling pathway. Whether these transcription factors are direct targets of miR-1 regulation remains to be seen. Notably, miR-1 predicted targets also include pro-ageing components of mTOR signalling such as *let-363/*mTOR and *raga-1/*RagA (targetscan.org), which also regulate the activity of these transcription factors. Hence the balance or timing of pro- and anti-ageing factors could also differentially influence the physiologic phenotype at a systemic level. Interestingly, VHA-13/ATP6VIA subunit has been shown to regulate mTOR lysosomal recruitment and activity (Chung et al., 2019) and mTOR signalling regulates the activity of TFEB at the lysosomal surface (Settembre et al., 2013). Upregulation of v-ATPase activity is also associated with the clearance of oxidized protein and rejuvenation of the *C. elegans* germline (Bohnert & Kenyon, 2017). Thus, in the future it will be interesting to further unravel the miR-1 molecular circuitry surrounding lysosomal function and proteostasis, and see whether miR-1 similarly regulates v-ATPase function in humans.

## Materials and Methods

### *C. elegans* strains and culture

*C. elegans* strains were maintained at 20°C on nematode growth medium (NGM) plates seeded with a lawn of *E. coli* strain OP50, unless noted otherwise.

N2 Bristol (wild-type) CGC

AA2508, *mir-1(gk276)* I

AA4575, *mir-1(dh1111)* I

AM140, *rmIs132[unc-54p::Q35::YFP]* I

AA4403, *mir-1(gk276); rmIs132[unc-54p::Q35::YFP]* I

AA4577, *mir-1(dh1111) rmIs132[unc-54p::Q35::YFP*] I

AA3275, *N2; dhEx965[mir-1p::mir-1, myo-2p::mCherry]*

AA4810, *mir-1(gk276)* I; *rmIs132[unc-54p::Q35::YFP]; dhEx965[mir-1p::mir-1, myo-2p::mCherry]*

AA4865, N2; *dhEx1206[myo3p::flag::HA::mCherry::vha-13cDNA::unc-54* 3’UTR*, myo-2p::GFP]*

AA4866, *mir-1(gk276)* I; *dhEx1206[myo-3p::flag::HA::mCherry::vha-13cDNA::unc-54* 3’UTR*, myo-2p::GFP]*

AA5067, N2; dh*Ex1207[myo-3p::flag::HA::mCherry::vha-13cDNA::vha-13 3’UTR, myo-2p::GFP]*

AA5068, *mir-1(gk276)* I*; dhEx1207[myo-3p::flag::HA::mCherry::vha-13cDNA::vha-13 3’UTR, myo-2p::GFP]*

PHX586, *vha-13(syb586[3xFlag::mNeonGreen::vha-13])* V

AA4813, *mir-1(gk276)* I; *vha-13(syb586[3xFlag::mNeonGreen::vha-13])* V

AA4850, *vha-13(syb586[3xFlag::mNeonGreen::vha-13])* V; *dhEx965[mir-1p::mir-1, myo-2p::mCherry]*

PHX587, *vha-13(syb587,syb504[3xFlag::mNeonGreen::vha-13* miR-1 BS1 mutated]) V. Also named in the text *vha-13* MUT1.

AA4184, *mir-1(gk276)* I; *vha-13(syb587,syb504[3xFlag::mNeonGreen::vha-13* miR-1 BS1 mutated]) V

PHX2180, *vha-13(syb2180,syb587,syb504[3xFlag::mNeonGreen::vha-13* miR-1 BS1,2 mutated]) V. Also named in the text *vha-13* MUT2.

AA5123, *mir-1(gk276)* I; *vha-13(syb2180,syb587,syb504[3xFlag::mNeonGreen::vha-13* miR-1 BS,2 mutated]) V

PHX1093, *vha-5(syb1093[3xFlag::mNeonGreen::vha-5])* IV

AA5069, *mir-1(gk276)* I; *vha-5(syb1093[3xFlag::mNeonGreen::vha-5])* IV

PHX809, *hlh-30(syb809[hlh-30::mNeonGreen])* IV

AA5195, *mir-1(gk276)* I; *hlh-30(syb809[hlh-30::mNeonGreen])* IV

### Molecular cloning

All restriction digest reactions were performed with enzymes provided by NEB according to the user’s manual. T4 DNA Ligase (NEB) was used for ligation reactions. Chemically competent DH5α *Escherichia coli* (LifeTechnologies) was used for transformation following the manufacturer’s instructions. QIAprep Miniprep or Midiprep Kits (Qiagen) were used for plasmid purification. Cloning was verified by PCR followed by gel electrophoresis and sequencing.

To make the rescuing *mir-1* transgene, primer pair "celmir1fwd2/rvs2" (Table S8) was used to insert the *mir-1* coding region into vector L3781 downstream of *gfp*. The *mir-1* promoter was then cloned 5’ to *gfp* using the primer pair "m1p fwd/rvs”(Table S8). To make muscle expressed *vha-13* constructs, the *vha-13* cDNA was amplified with Kpn1 overhangs using primers vha-13 fwd/rvs (Table S8) and cloned into vector pDESTR4-R3 to give *myo3p::flag::HA::mCherry::vha-13cDNA::unc-54* 3’UTR. The *unc-54* 3’UTR was excised with Not1/BglI and replaced with the *vha-13* 3’UTR using primers vha-13U fwd/rvs (Table S8) to generate *myo3p::flag::HA::mCherry::vha-13cDNA::vha-13 3’UTR.*

### Generation of transgenic worm strains

Transgenic worms containing extrachromosomal arrays were generated by microinjection. To generate the ***myo3p::flag::HA::mCherry::vha-13cDNA::unc-54* 3’UTR** strain, a mix of *myo3p::flag::HA::mCherry::vha-13cDNA::unc-54* 3’UTR DNA (40ng/ μl), *myo-2p::gfp* co-transformation marker (5 ng/ μl plasmid pPD122.11) and fill DNA (TOPO empty vector, 55 ng/ μl) was injected into young N2 adults using an Axio Imager Z1 microscope (Zeiss) with a manual micromanipulator (Narishige) connected to a microinjector (FemtoJet 4x, Eppendorf). We obtained the strain AA4865: N2*; dhEx1206[myo3p::flag::HA::mCherry::vha-13cDNA::unc-54 3‘UTR, myo-2p::gfp]*, which was then used to cross the transgene into other genetic backgrounds. A similar strategy was used to create AA5067, N2; dh*Ex1207[myo3p::flag::HA::mCherry::vha-13cDNA::vha-13 3’UTR, myo-2p::GFP].*

The ***mir-1* transgene** strain was generated by injection mix containing *mir-1p::mir-1* plasmid, *myo-2p::mCherry* co-transformation marker (5 ng/ μl plasmid pPD122.11) and fill DNA (L3781 empty vector). The resultant strain AA3275 (*N2; dhEx965[mir-1p::mir-1, myo-2p::mCherry]*) was then used to cross the *dhEx965* transgene to *mir-1(gk276); rmIs132[p(unc-54) Q35::YFP]* to give *mir-1(gk276); rmIs132[p(unc-54) Q35::YFP]; dhEx965[mir-1p::mir-1, myo-2p::mCherry]*, AA4810. For transgenic worm strains, non-transgenic worms of the same strain were used as controls.

The ***mir-1* deletion allele *dh1111*** was generated using CRISPR/Cas9 mutagenesis. We designed CRISPR guides using the EnGen sgRNA Designer (https://sgrna.neb.com/#!/sgrna), synthesized guides with the Engen sgRNA synthesis kit and analysed them by gel electrophoresis and tape station. We injected worms with an injection mix containing Cas9 EnGen (NEB), 4 sgRNAs against *mir-1* (AAGAAGTATGTAGAACGGGG, GTAAAGAAGTATGTAGAACG, TATAGAGTAGAATTGAATCT, ATATAGAGTAGAATTGAATC), one sgRNA against *dpy-10* (CGCTACCATAGGCACCACG), KCl, Hepes pH 7.4 and water. Prior to injection we incubated the mixture for 10 minutes at 37°C to allow activation of Cas9. Following injection, we singled out worms with Dpy phenotype in the F1 generation and genotyped them for *mir-1* deletion using *mir-1* genotyping primers (Supplemental Table 8). We sequenced the PCR products of candidate worms with Sanger sequencing and verified that the deletions resulted in loss of *mir-1* expression using Taqman-based quantification of mature miRNA levels.

### Endogenous fluorescently tagged strains

were generated by tagging *vha-5* or *vha-13* with 3xFLAG-mNeonGreen tag at the N-terminus using CRISPR–Cas9 (SunyBiotech). The *vha-13* 3‘UTR mutants BS1 and BS2 were generated using CRISPR–Cas9, by further mutating one or both putative miR-1 binding sites in the 3’UTR of the endogenous FLAG-mNeonGreen-tagged *vha-13 gene* (SunyBiotech). PHX809 *hlh-30::mNeonGreen* endogenously tagged *hlh-30* was generated by placing *mNeonGreen* at the C-terminus using CRISPR–Cas9 (SunyBiotech).

### Determination of progeny number

Single worms were maintained on an NGM agar plate and transferred every day until the reproductive period was complete. The number of F1 progeny per individual worm was counted at the L4 or young adult stage. Experiments were repeated at least three times.

### Quantitation of polyQ aggregates

Whole-worm images of day 4 adults were taken with an Axiocam 506 mono (Zeiss) camera using the 5x objective of a Zeiss Axio Imager Z1 microscope (20ms exposure). Q35 aggregate numbers were evaluated from photos at 163% magnification. Genotypes of the samples were blinded during the counting. Aggregates were defined as discrete structures or puncta above background.

### Motility, thrashing and pumping behaviour

To analyse worm motility by the circle test, 20 to 25 worms were placed in the centre of a 6 cm agar plate with bacteria, marked with 1 cm circle on the bottom. After a defined time period (1 or 30 minutes) the number of worms that left the circle were counted. To determine thrashing rate, individual worms were transferred to M9 buffer and the number of body bends in a 20 to 30 second interval was scored. Pharyngeal pumping rates were measured by counting grinder contraction in the terminal bulb over 20 to 30 seconds. Genotypes were blinded during all experiments.

### CeleST Assay

The *C. elegans* Swim Test (CeleST) assay was used to assess animal motility while swimming; assays were conducted as described previously (Ibanez-Ventoso, Herrera, Chen, Motto, & Driscoll, 2016; Restif et al., 2014). Four or five animals were picked and placed into the swimming arena, which was a 50 μl aliquot of M9 Buffer inside a 10 mm pre-printed ring on the surface of a glass microscope slide (Thermo Fisher Scientific). Images of the animals within the swim arena were acquired with a LEICA MDG41 microscope at 1x magnification with a Leica DFC3000G camera. Image sequences of 30 seconds in duration were captured at a rate of ~16 frames per second. The established CeleST software was used to process the image sequences and extract 8 measures that are descriptive of *C. elegans* swim motility. As not all images in a sequence are always successfully processed by the CeleST software, animals for which fewer than 80% of frames were valid were excluded from the analysis. For each measure, single measurements that did not fit within the range of normally observed values were deemed outliers and also excluded from the analysis. Unpaired Student’s t-test was used to test for statistical significance between two strains at adult day 4 and day 8.

### Life span and heat stress experiments

Life span experiments were performed as described (Gerisch et al., 2007). Day 0 corresponds to the L4 stage. Life spans were determined by scoring a population of 100 to 120 worms per genotype every day or every other day. Worms that exploded, had internal hatch or left the plate were censored. Heat stress experiments were performed at 35°C (Gerisch et al., 2007). Day 1 adult worms were transferred onto pre-heated (35°C) plates 6 cm plates seeded with OP50. Worms were kept at 35°C and scored every hour for live versus dead. Experiments were repeated at least three times. Data were analysed with Microsoft Excel 16.12 and GraphPad Prism 7 Software. *P*-values were calculated using the Log-rank (Mantel-Cox) test to compare two independent populations.

### RNAi Treatment

Worms were grown from L4 onwards unless mentioned otherwise on *E. coli* HT115 (DE3) bacteria expressing dsRNA of the target gene under the control of an IPTG-inducible promotor. RNAi colonies were grown overnight at 37°C in Luria Broth with 50 μg/ml Ampicillin and 10 μg/ml tetracycline. The cultures were spun down at 4,000 rpm at 4°C for 10 minutes. 500 μl of one-fold concentrated culture was seeded onto NGM plates containing 1 M isopropyl β-D-1-thiogalactopyranoside (IPTG) to induce dsRNA expression. RNAi clones were selected from the Ahringer or Vidal library (Kamath & Ahringer, 2003; Rual et al., 2004). Clone identity was confirmed by sequencing. Bacteria containing luciferase were used as control.

### RNAi screen for *mir-1* suppressors

*mir-1(gk276)* Q35 worms were grown from L4 on RNAi of the candidates identified in the bioinformatic or proteomic screen. Worms were maintained in the presence of FUDR till day 8 of adulthood. The circle test was performed as described above. The number of worms that left the circle was determined after 1 minute. Selected candidates identified in the circle test were examined in the body bending assay (see above). The effect of selected candidate RNAi clones was additionally tested on polyQ35 in the wild-type background as counter screen. For this experiment worms were grown till day 5 of adulthood. Motility assays were performed on day 5 of adulthood because Q35 wild-type worms were less motile than the *mir-1(gk276)* Q35 and were largely paralyzed by day 8.

### RNA extraction and real-time qPCR analysis

Worm populations (ca. 500 animals) were harvested on day 4 of adulthood and washed twice in cold M9. Worm pellets were taken up in 700 μl QIAzol reagent (Qiagen) and snap frozen in liquid nitrogen. Samples were subjected to 4 freeze/thaw cycles and homogenized with 1.0 mm zirconia/silica beads (Fisher Scientific) in a Tissue Lyser LT (Qiagen) for 15 min at full speed. After homogenization, 600μl supernatant was transferred to fresh tubes and 120 μl chloroform were added to each tube. Components were mixed by vortexing and incubated for 2 min at room temperature. After 15 min centrifugation at 12,000 *g* 4°C, the aqueous phase was collected for total RNA extraction using the RNeasy or miRNeasy Mini Kit (Qiagen) according to manufacturer’s instructions. RNA quantity and quality were determined on a NanoDrop 2000c (PeqLab) and cDNA was prepared using the iScript cDNA Synthesis Kit (BioRad). To quantify RNA expression, we used the Power SYBR Green Master Mix (Applied Biosystems) on a ViiA 7 Real-Time PCR system (Applied Biosystems). Four technical replicates were pipetted on a 384-well plate using the JANUS automated workstation (PerkinElmer). Expression of target RNA was calculated from comparative CT values, normalized to *ama-1* or *cdc-42* as internal controls using the corresponding ViiA7 software. All unpublished primers were validated by determination of their standard curves and melting properties. For quantification of *mir-1* we used Taqman probes from LifeTechnologies/ThermoFisher Scientific (Assay ID 000385) and normalized to expression of U18 as measured by Taqman probes from LifeTechnologies/ThermoFisher Scientific (Assay ID 001764). We used N=4 independent biological replicates, and four technical replicates for every biological sample (for primers, refer to Supplementary Table 8).

### Western blot analysis

For Western blot analysis synchronized young adult or gravid day 1 adult worms were picked into Eppendorf tubes containing M9, snap frozen in liquid nitrogen and lysed in 4x SDS sample buffer (Thermo Fisher) containing 50 mM DTT. After boiling and sonication, equal volumes were subjected to reducing SDS-PAGE and transferred to nitrocellulose membranes. The membranes were then blocked for two hours at room temperature in 5% milk in Tris-buffered Saline and Tween20 (TBST) and probed with the primary antibodies in TBST with 5% milk overnight at 4 °C. Specific secondary antibodies (mouse or rabbit) were used at a concentration of 1:5000 in TBST with 5% milk at room temperature for 2 hours. The membranes were developed with Western Lightening Plus- Enhanced Chemiluminescence Substrate (PerkinElmer). Bands were detected on a ChemiDoc MP Imaging System (BioRad) and the intensity quantified using the corresponding Image Lab software (BioRad). The following antibodies were used: anti-GFP (JL-8 Living Colors, mouse monoclonal), anti-Histone H3 (ab1791 Abcam, rabbit polyclonal), anti-α-Tubulin (T6199 Sigma, mouse monoclonal), anti-FLAG (F1804 Sigma, mouse monoclonal), anti-mouse IgG (G-21040 Invitrogen, goat polyclonal coupled with horseradish peroxidase), anti-rabbit IgG (G-21234 Invitrogen, goat polyclonal coupled with horseradish peroxidase).

### Proteomic analysis

For **sample collection and preparation**, day 1 N2 and *mir-1(gk276)* worms were synchronized by egg laying, and *n* ≥ 5000 worms per genotype were collected in M9. Samples were washed three times in M9 and directly frozen in liquid nitrogen and stored at −80 °C. 5 independent biological replicates of each genotype were collected for further analyses. For **protein extraction** samples were thawed and boiled in lysis buffer (100 mM Tris-HCl, 6 M guanidinium chloride, 10 mM Tris(2-carboxyethyl)phosphine hydrochloride, 40 mM 2-Chloroacetamide) for 10 min, lysed at high performance with a Bioruptor Plus sonicator (Diagenode) using 10 cycles of 30 s sonication intervals. The samples were then boiled again, centrifuged at 20000*g* for 20 min, and diluted 1:10 in 20 mM Tris pH 8.3 / 10% acetonitrile (ACN). Protein concentration was measured using BCA Protein Assays (Thermo Fisher). Samples were then digested overnight with rLys-C (Promega), the peptides were cleaned on a Supelco Visiprep SPE Vacuum Manifold (Sigma) using OASIS HLB Extraction cartridges (Waters). The columns were conditioned twice with Methanol, equilibrated twice with 0.1% formic acid, loaded with the sample, washed three times with 0.1% formic acid and the peptides eluted with 60% ACN / 0.1% formic acid. The samples were dried at 30°C for roughly 4 h in a Concentrator (Eppendorf) set for volatile aqueous substances. The dried peptides were taken up in 0.1% formic acid and the samples were analysed by the Max Planck Proteomic Core facility. **Mass spectrometry** data acquisition, computational proteomic analysis and differential expression analysis were performed as described (Tharyan et al., 2020). Upon inspection of the numbers of quantified proteins and the raw proteomic data, two replicates of N2 were excluded from further analysis.

### Microscopy and expression analysis

For confocal microscopy *vha-13*, *vha-5 and hlh-30* endogenously tagged with mNeonGreen were synchronized via egg and maintained at 25°C to induce mild stress. *hlh-30* worms were imaged as day 4 adults, while *vha-13* and *vha-5* worms as L4s, because of high mNeonGreen background expression at day 4 adulthood. Worms were anaesthetized with 40 μM sodium azide, mounted on slide with 2% agar pad and imaged with a Leica TCS SP8 microscope equipped with HC PL APO CS2 63X/1.40 Oil and white light laser. Images were analysed using photoshop or Fiji software. To quantitate acidic lysosomes, worms grown at 25°C were incubated 48 hours prior to imaging with 2 μM LysoTracker Red DND-99 (Life Technologies). Worms were imaged as day 4 adults as described above. Number of puncta in a predefined squared area in the intestine (approximately spanning the second to fourth gut cell) were counted.

### Statistical Analysis

The statistical tests performed in this study are indicated in Figure legends and in the method detail section. Data are represented as mean ± SEM or as individual data points, as stated in the Figure legends. Number of replicates and animals for each experiment are enclosed in their respective Figure legends.

### Data and Software Availability

Accession Code: The mass spectrometry proteomics data have been deposited to the ProteomeXchange Consortium via the PRIDE partner repository (Perez-Riverol et al., 2019) with the dataset identifier PXD023544.

## Acknowledgements

AA would like to thank the MPI-AGE proteomics and imaging cores for services, the Caenorhabditis Genetics Center (CGC, University of Minnesota) for worm strains, the Bundesministerium für Bildung und Forschung for Sybacol funding, the Deutscher Akademischer Austauschdienst for funding, and the Max Planck Gesellschaft for core institutional support. MAM would like to thank Fundação de Amparo à Pesquisa do Estado de São Paulo (FAPESP) (Grant number 2019/25958-9) and Coordenação de Aperfeiçoamento de Pessoal de Nível Superior CAPES (Grant number 88881.143924/2017-01) for funding.

## Contributions

IS, BG, KK, RL, JH, MSD, YS, AA designed and performed experiments IS, BG, SV, OS, AA wrote the paper

## Supplementary data

**Supplementary Figure 1.**
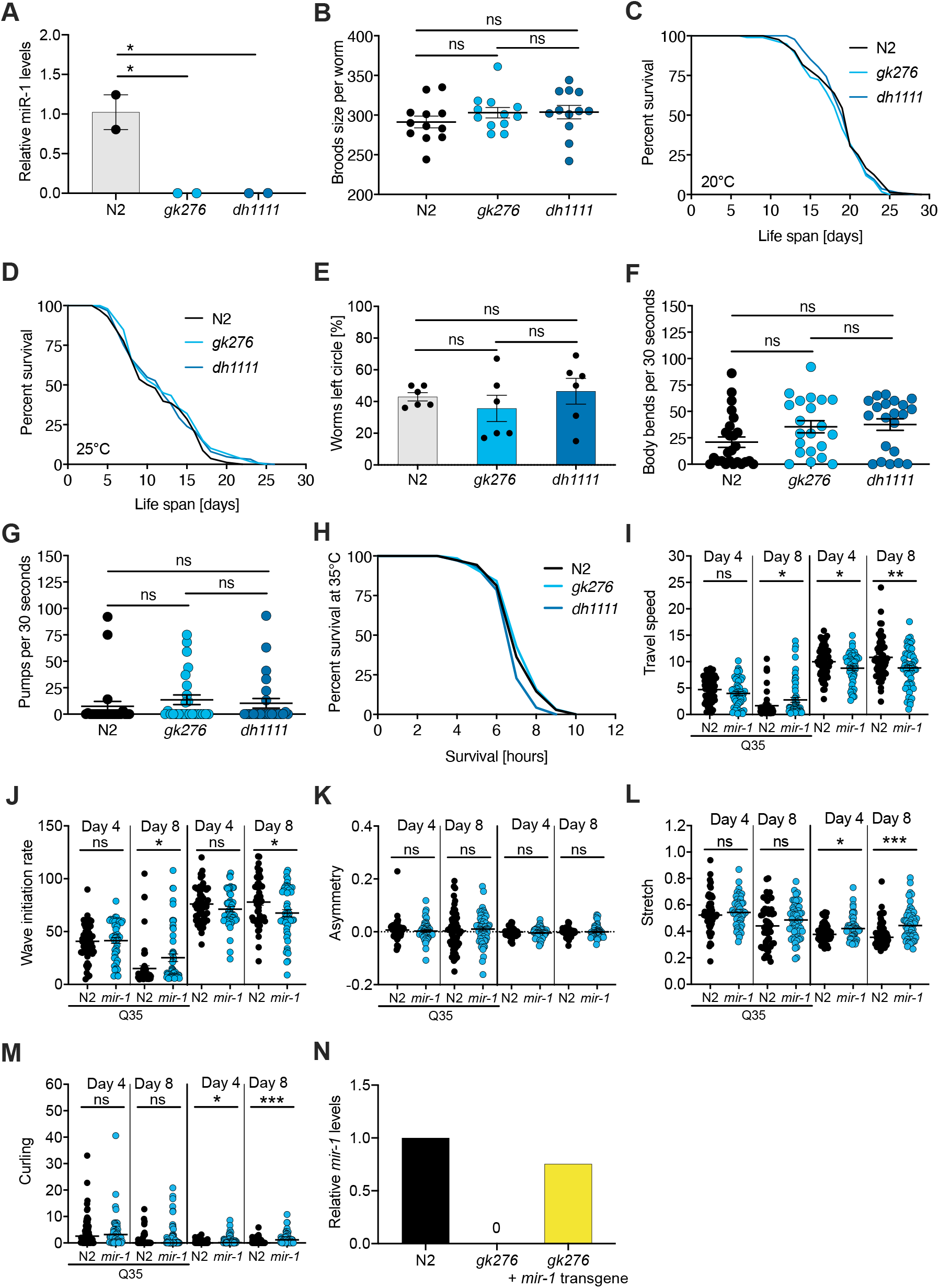
(linked to Figure 1) **(A)** TaqMan qPCR measuring mature *mir-1* levels show that the deletion alleles do not express *mir-1* and are null, N=2 BR. mean ± SEM, one way ANOVA, *, p<0.05. **(B)** Brood size of *mir-1(gk276)* and *mir-1(dh1111)* compared to wild-type (N2). Each dot represents the brood size of one worm, mean ± SEM, one way ANOVA, ns, not significant. **(C)** and **(D)** Lifespan experiments performed at 20°C and 25°C for two *mir-1* deletion alleles compared to wild-type. N=3. Not significant (Supplemental Table 1). **(E)** and **(F)** Motility of wild-type and *mir-1(gk276)* mutant animals on day 14 of adulthood, measured by circle test (E) and thrashing in liquid (F). One representative experiment of each N=3 is shown. Mean ± SEM, one way ANOVA, ns, not significant. **(G)** Pharyngeal pumping rate measured on day 14 in two *mir-1* alleles compared to wild-type. N=3, one representative experiment is shown. Mean ± SEM, one way ANOVA, ns, not significant. **(H)** Heat stress survival at 35°C of day 1 *mir-1(gk276)* and *mir-1 (dh1111)* mutants compared to wild-type. One representative experiment is shown, N=3, t-test: N2 *vs. gk276*: p=0.20. N2 *vs. dh1111*: p=0.17. **(I)** to **(M)** CeleST locomotion behaviour analyses of wild-type and *mir-1(gk276)* mutant animals with or without *unc-54p:Q35:YFP*, at day 4 and day 8 of adulthood. Each dot represents one animal. **(I)** travel speed **(J)** wave initiation rate **(K)** asymmetry **(L)** stretch, and **(M)** curling. t-test, *, p<0.05, **, p<0.01, ***, p<0.001, ns, not significant. **(N)** Bar graph showing the expression level of mature miR-1 microRNA in the indicated genotypes. N=1 biological replicate, 4 technical replicates.

**Supplementary Figure 2.**
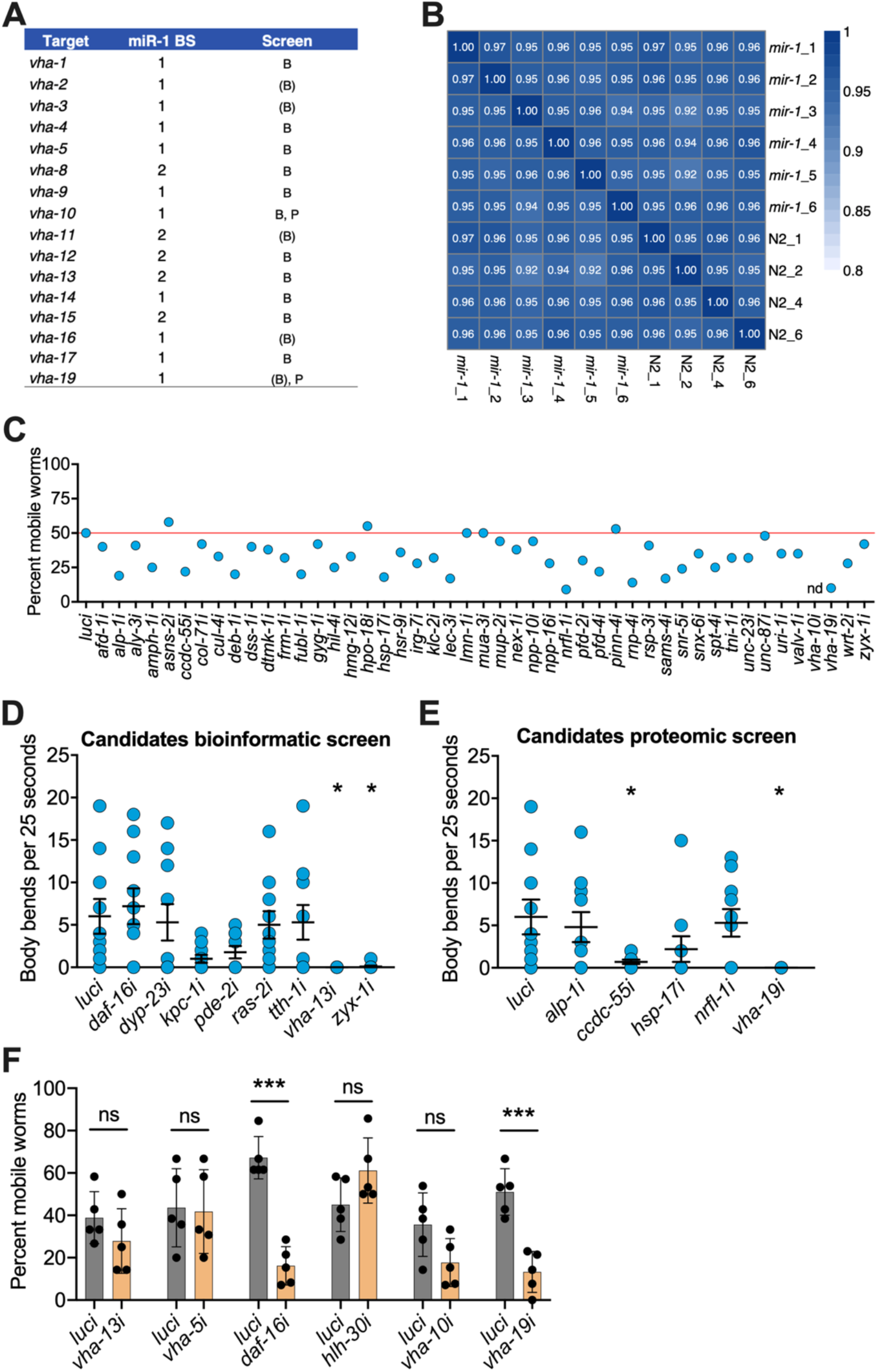
(linked to Figure 2) **(A)** v-ATPase subunits containing one or two predicted miR-1 binding sides (BS) in their 3’UTR. v-ATPase genes identified in the bioinformatic screen by all three databases are labelled „B“, by 2 databases are labelled „(B)“ and by the proteomic screen are labelled „P“. **(B)** Correlation plot of the biological replicates from proteomics showing the similarity of WT (N2) and *mir-1* genotypes. **(C)** RNAi screen for genes required for *mir-1(gk276)*;Q35 motility in the circle test, using upregulated genes from the proteomic analysis (Supplemental Tables 5 and 6). Each dot represents the percentage of worms that left the circle. Red line, percent luciferase controls that left the circle. Nd, not determined. Validation of selected candidates identified from bioinformatic **(D)** and **(E)** proteomic screens by thrashing assay on day 8 of adulthood. *mir-1(gk276)*;Q35 worms were grown on the corresponding RNAi from L4 onwards. Each dot represents one animal, 15 worms per RNAi. Mean ± SEM, one way ANOVA, only significant values are labelled: *, p<0.05. **(F)** Motility of worms in circle test of indicated RNAi on N2;Q35 worms. Motility was measured on day 5 of adulthood, each dot represents the percentage of worms that left the circle. Mean ± SEM of one representative experiment, N=3, one way ANOVA, ***, p<0.001, ns, not significant.

**Supplementary Figure 3.**
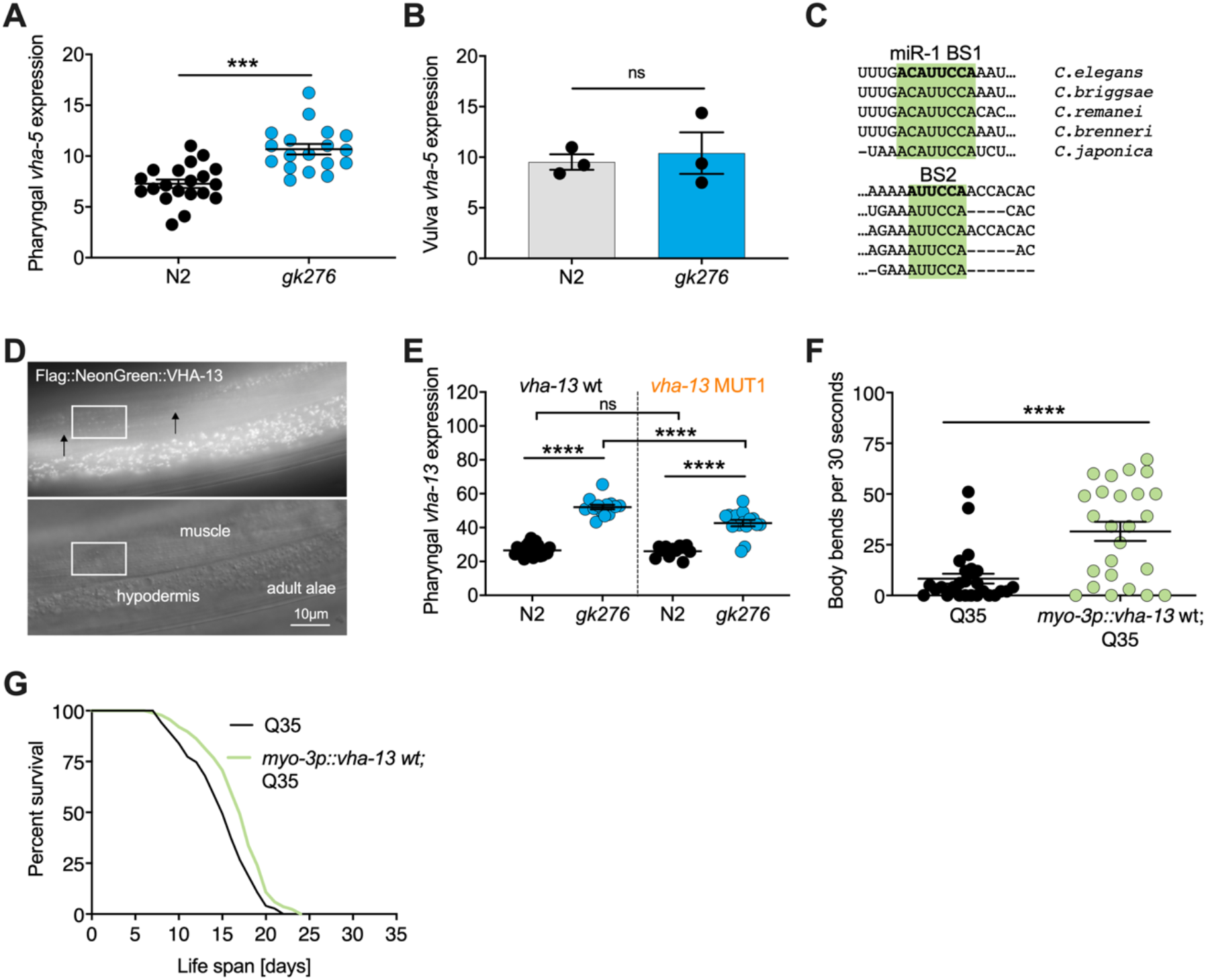
(linked to Figure 3) **(A)** Quantification of pharyngeal fluorescent intensity of endogenous *3xFlag:mNeon Green:vha-5* expression in indicated genotypes in late L4s maintained at 25°C. One representative experiment of N=3, mean ± SEM, t-test, ***, p<0.001. **(B)** Quantification of fluorescent intensity of *3xFlag:mNeonGreen:vha-5* expression in the vulva of indicated genotypes. Mean ± SEM of N=3, t-test, ns, not significant. **(C)** Schematic showing the conservation of miR-1 binding sites (BS) in the *vha-13* 3’UTR of different nematode species. **(D)** Images showing expression of endogenously tagged *3xFlag:mNeonGreen:vha-13* in muscle dense body and hypodermis. Arrows indicate individual dense bodies. Rectangle highlights muscle section shown in Figure 3B. **(E)** Quantification of fluorescence intensity in the isthmus of L4 larvae with endogenously tagged *3xFlag:mNeonGreen:vha-13* with *vha-13* MUT1 3’ UTR in relation to *vha-13* wt 3’UTR, in N2 and *mir-1(gk276)* mutant backgrounds using confocal microscopy. Mean ± SEM of one representative experiment. N=2, one way ANOVA, ****, p<0.00001, ns not significant. **(F)** Thrashing assay of *unc-54p:Q35:YFP* (Q35) worms expressing *vha-13* in body wall muscle (*myo3p*:*vha-13* wt;Q35) or non-transgenic Q35 animals. 25 worms per genotype, mean ± SEM of one representative experiment, N=4, t-test, ****, p<0.0001. **(G)** Life span of Q35 worms overexpressing *vha-13* in the body wall muscle (*myo3p*:*vha-13* wt;Q35) and Q35 non-transgenic segregants of the same strain. One experiment of N=4. Life span effects of two experiments were significant (Supplemental Table 1). t-test: 0.0017.

**Supplementary Figure 4.**
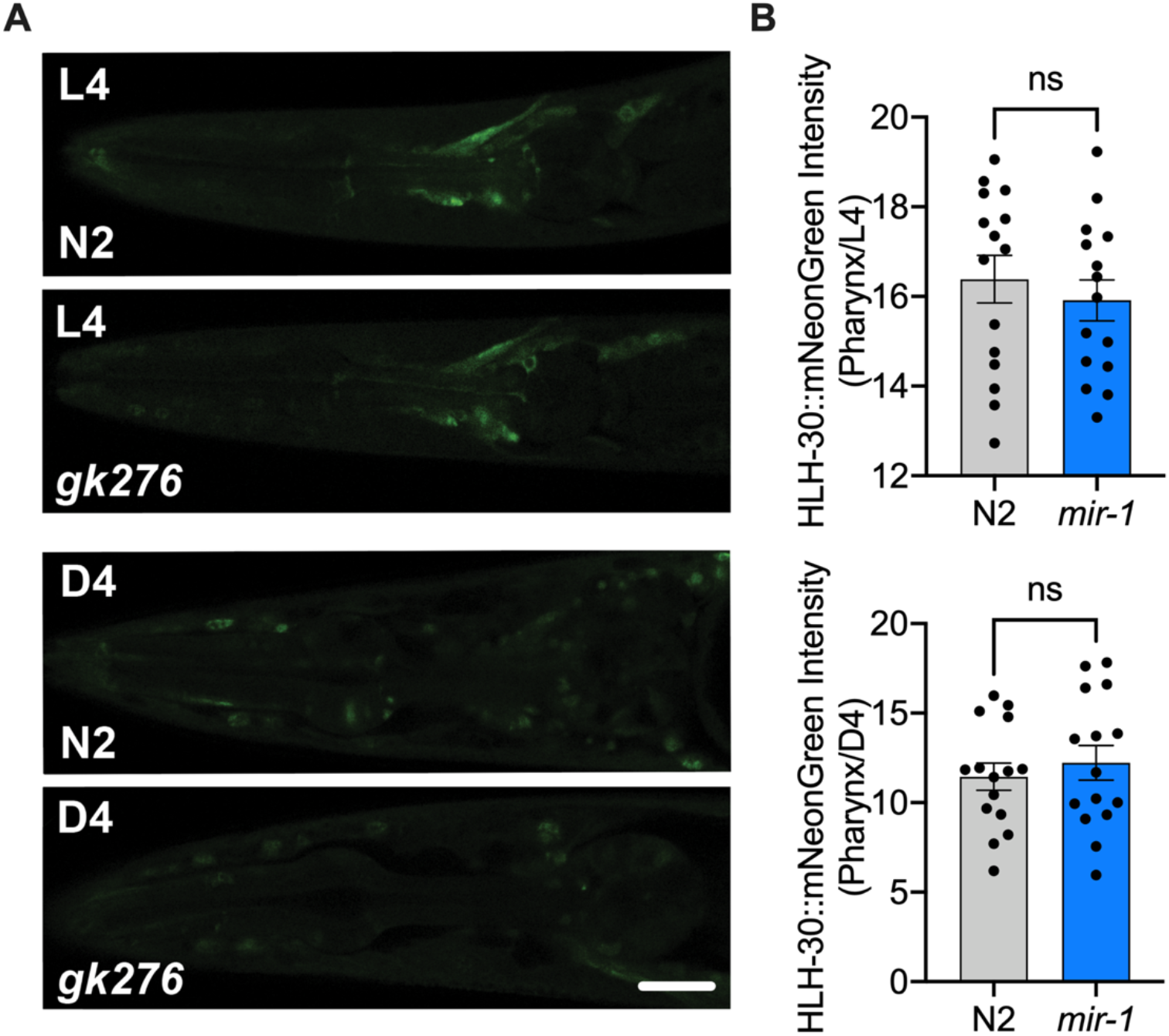
(linked to Figure 4) **(A)** Fluorescent image comparing HLH-30:mNeonGreen intensity in the pharynx of the *mir-1(gk276)* and WT (N2) backgrounds in L4s (top) or day 4 adults (D4, bottom). Scale bar 20 μm **(B)** Quantitation of nuclear localization in **(A)**, dots represent individual animals. N=2 biological replicates for L4 and N=1 for day 4 adults, t-test, ns: not significant.

---

Supplemental Table 1: Lifespan and heat stress data.

Supplemental Table 2: Q35 aggregate and behavior data.

Supplemental Table 3: CeleSt data.

Supplemental Table 4: Bioinformatic screen data.

Supplemental Table 5: RNAi motility screen.

Supplemental Table 6: Proteomic analysis of differentially regulated proteins in *mir-1 vs.* wildtype.

Supplemental Table 7: Microscopy data.

Supplemental Table 8: Primers.

